# Toward temporally calibrated biomarkers of heat stress in free-living songbirds

**DOI:** 10.64898/2026.06.16.732729

**Authors:** Isaac Miller-Crews, Elizabeth P. Derryberry, Kimberly A. Rosvall

## Abstract

As heatwaves increase in intensity and frequency, more birds are exposed to sublethal heat, which can affect many elements of the phenotype, from growth to cognition to reproduction. These widespread performance-related effects of heat, coupled with the rapid declines seen in many bird populations in recent decades, underscore the urgency of detecting recent heat exposure and its downstream physiological effects in the wild. To develop minimally invasive biomarkers of past heat, we experimentally elevated nest temperatures for free-living nestling Tree Swallows (*Tachycineta bicolor*) for four hours on their twelfth day of life. Twenty-four hours later, we returned to collect a small blood sample and quantify carryover effects of prior sublethal heat on the blood transcriptome. By comparing these carryover effects to those that occur in the immediate aftermath of heat, we identify biomarkers of heat that reflect distinct and time-dependent processes. Candidate biomarkers include four upregulated genes with connections to stress and disease (*LAMA3, ATP1B1, RASGEF1A, TMEM181*) and two additional down-regulated genes. By incorporating the sex of each nestling into our analyses, we also unveiled marked sexual dimorphism in the blood transcriptome, even among autosomal genes and including pathways that imply inherent sex differences in heat tolerance. When these sex differences are controlled, we see that the sexes respond to heat with overwhelming similarity, further grounding the utility of our suggested transcriptomic biomarkers. Though these biomarkers will require further validation to be used across bird species, our collective results uncover temporally calibrated targets can be measured with just one drop of blood, improving our understanding of climate impacts on wild birds.

**Lay summary:**

- As global temperatures rise, many birds experience bouts of heat stress, but we do not have simple biomarkers that reliably reflect this past exposure in the wild.
- We tested whether a small blood sample could reveal recent heat stress through changes in gene activity.
- Our experiment exposed nestling Tree Swallows to a non-lethal heat stressor and measured how their gene activity changed during and after the heat event.
- Some genes reacted quickly but returned to normal within a day, while others showed longer-lasting effects.
- Males and females responded to heat in similar ways, even though their baseline gene activity differed substantially.
- Six genes responded consistently across the sexes and in relation to temperature, making them promising biomarkers of past heat.
- These results can help scientists better track heat exposure in wild birds and improve predictions on how populations respond to continued climate change.

## Introduction

As global temperatures rise, heatwaves have become more common and more intense (Fischer et al., 2021), shifting the selective environment for organisms across the globe (Buckley and Huey, 2016). At the same time, bird populations are declining (Rosenberg et al., 2019), and heat is one key driver of these losses (Halupka et al., 2023; Molenaar et al., 2026). Elevated temperatures have triggered mass mortality events in birds and other small endotherms (Kim and Stephen, 2018; McKechnie and Wolf, 2010; McKechnie and Wolf, 2019; Riddell et al., 2021; Welbergen et al., 2008), yet it is the sublethal effects of heat that are thought to be particularly widespread and impactful (Angilletta, 2009; Conradie et al., 2019; Stillman, 2019). For example, among adult birds, common spring and summer temperatures can impair cognitive performance, interfere with mate choice, and reduce critical mating traits like the singing of songs or production of sperm (Coomes et al., 2019; Danner et al., 2021; Hurley et al., 2018; Leith et al., 2022; Lipshutz et al., 2022; Soravia et al., 2021; Soravia et al., 2023). Likewise, in some studies, heat-exposed nestlings or juveniles invest more in thermoregulatory responses that trade-off with other fitness-critical traits like begging for food (Derryberry et al., in press) or somatic growth (Andreasson et al., 2018; Andrew et al., 2017; Corregidor-Castro and Jones, 2021). Some of these early life heat effects may carry-over to adulthood, shaping later physiology, cognition, and even ageing (Hoffman et al., 2024; Hoffman et al., 2026; Soravia et al., 2024; Stier et al., 2024). Despite growing recognition of these diverse heat impacts on breeding birds and their offspring, our ability to detect and interpret heat exposure in the wild remains limited (Morosinotto et al., 2025).

Biomarkers can help with this challenge because they capture some element of the natural biological response to an environmental stimulus. There are many types of biomarkers (Califf, 2018), but the most useful subtypes are *monitoring* or *response* biomarkers, which assess evidence of exposure to some biotic or abiotic agent. For example, aspects of stress physiology are increasingly measured in wild plants and animals to assess the impacts of global climate change (Beaulieu and Costantini, 2014; Costantini et al., 2025; Dantzer et al., 2014; Madliger et al., 2018; Schönbeck et al., 2023). Historically, most of this work focused on terrestrial mammals (Madliger et al., 2018) or aquatic ectotherms (Fangue et al., 2006; Kenkel and Matz, 2016; Li et al., 2019; Parkinson et al., 2020). Comparable data on the physiology of avian heat responses has lagged behind (McKechnie and Wolf, 2019; Nord and Giroud, 2020), though this knowledge gap is narrowing (Nord et al., 2026). Early insights on heat-related biomarkers in birds focused on agricultural contexts (Etches et al., 2008; Sejian et al., 2018) and certain heat shock proteins (HSPs) that elevate in poultry blood or feathers in the aftermath of heat (Dionello et al., 2001; Greene et al., 2019; Kang and Shim, 2021; Xie et al., 2014). As research on more wild species has emerged, we now know that HSP mRNA abundance can spike in the blood within four hours of sublethal heat (Woodruff et al., 2025a), even in altricial nestlings as young as six days post-hatch (Derryberry et al., in press). Mirroring findings across many vertebrate taxa (Dantzer et al., 2014), heat may trigger glucocorticoid secretion into the blood within minutes (Mentesana and Hau, 2022), with carryover effects embedded in feathers or fecal samples (Fairhurst et al., 2012; Moagi et al., 2021). Other potential biomarkers of heat include elements of oxidative damage (Beaulieu and Costantini, 2014; Ton et al., 2021), telomere shortening (Furic et al., 2026; Sohn et al., 2012), or even behavioral adjustments (Rosvall and Derryberry, in press), underscoring the broad suite of potential metrics that may reflect an organism’s recent exposure to sublethal heat.

Adding further complexity, many stress-reactive phenotypes exhibit an initial stress response that differs from its later post-stress condition, meaning that temporal dynamics are an important component of biomarker validation. For instance, with HSPs, *Acropora* corals have elevated abundance of some *HSP70* and *HSP90* transcripts at 24 hours after a heat event but not at 48 hours (Javid et al., 2025). In nestling Tree Swallows (*Tachycineta bicolor*) *HSP90AA1* mRNA in the blood is elevated immediately after a four-hour heat challenge; this is the most heat reactive HSP in Tree Swallow blood (Woodruff et al., 2025a), but those levels return to those of controls 24 hours later (Derryberry et al., in press). With glucocorticoids (Romero et al., 2009), an acute stressor tends to increase blood levels of cortisol or corticosterone, yet chronic stress can dampen an organism’s ability to mount an acute response (McEwen and Wingfield, 2003) and delay the organism’s return to baseline after a stressor (Gormally and Romero, 2018), with mixed evidence of either elevated (Lowrance et al., 2016) or diminished (Cyr et al., 2007) baseline glucocorticoid levels. Transcriptomic profiling can uncover new candidates beyond these classically considered stress markers because, for example, it can document all expressed genes that change in the blood in response to some stimulus (Louder et al., 2020; Meitern et al., 2014). However, the transcriptome is temporally dynamic (Rittschof and Hughes, 2018; Saul et al., 2019), suggesting that the genes whose transcription is most affected during, or immediately after, a heat challenge may be fundamentally different from those whose altered expression endures beyond the stressor itself, potentially providing a stronger signal of the biological embedding of past heat exposure. This hypothesis finds some support in domestic poultry (Xie et al., 2014), corals (Savary et al., 2021), and beyond (Ali et al., 2025; Jovic et al., 2019), but it remains to be tested in wild birds for whom we deeply need more information on individual, population, and species variation in the potentially lasting effects of heat.

Here, we devised a manipulate-and-sample-later design, experimentally elevating nest temperatures for free-living nestlings and measuring carryover effects of heat in the blood 24 hours later (see ***Figure 1A***). Specifically, we generated an intense but sublethal heat challenge for 12-day-old Tree Swallows confined to their nesting cavity, replicating our previous work that adds air-activated heat packs to the nestbox for four hours. In past iterations of this experiment, we have seen increases in panting and other thermoregulatory behaviors, alongside a spike in blood gene expression for many HSPs and other damage-repair processes at the end of the four-hour challenge (Woodruff et al., 2025a). Leveraging this past transcriptomic analysis with new blood-derived RNA-seq results collected 24-hours after heat, we aim to identify candidate biomarkers that can be used to assess past exposure to sublethal heat. To this end, we evaluated genes whose expression reflects only an initial reaction to heat, only an organism’s post-heat exposure condition, or some combination of the two. To refine these biomarkers for future application, we accounted for sex-specific variation while also distinguishing between biomarkers that capture discrete responses to heat versus those that encode graded information about thermal conditions within the nest. In doing so, we develop a minimally invasive and temporally calibrated diagnostic framework for detecting recent environmental heat stress in wild songbird populations.

**Figure 1.**
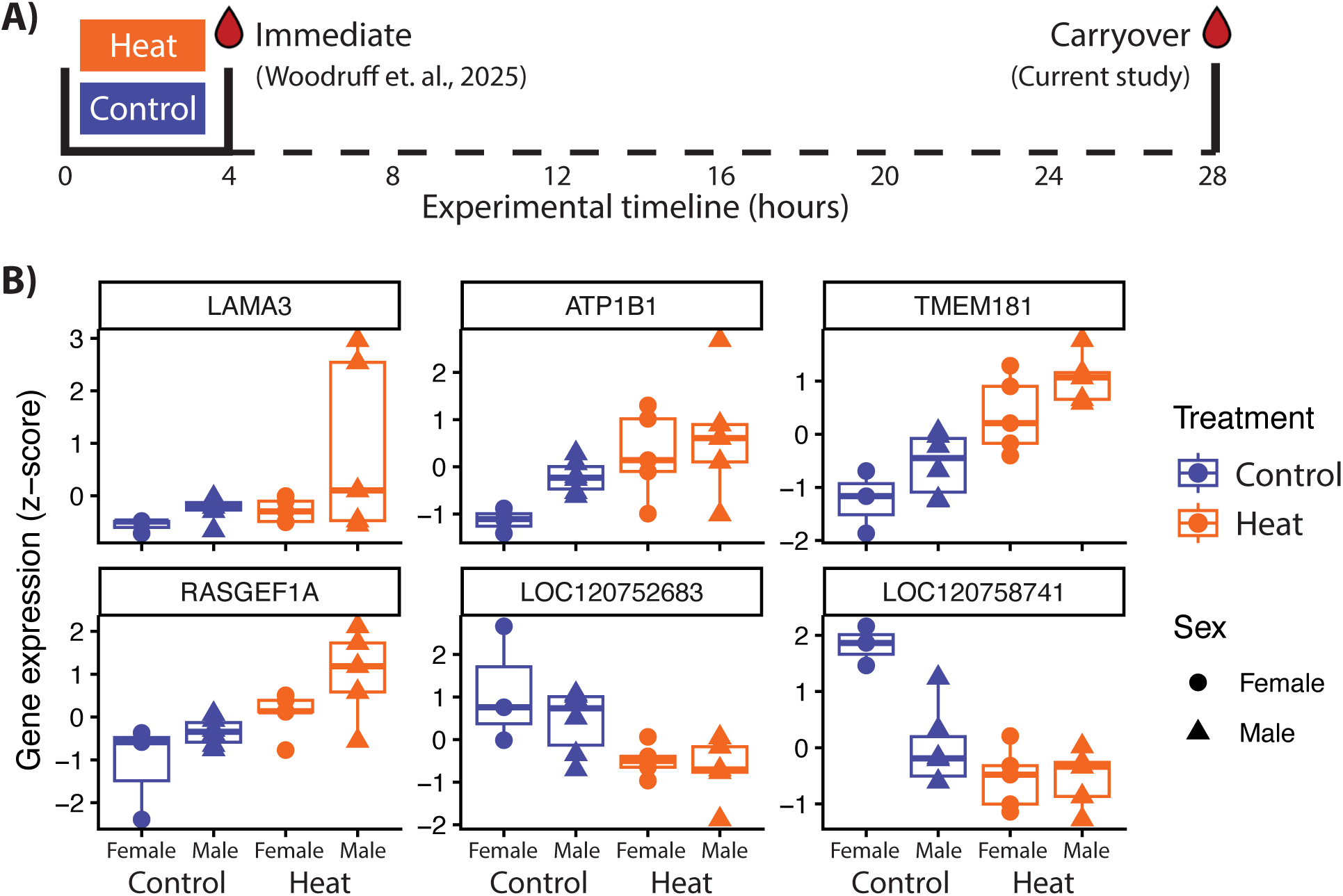
**A) Experimental design.** To understand carryover effects of past heat, this current study manipulated temperature for four hours in the nestbox, and then sampled 12-day-old nestlings 24 hours later (“Carryover”). In a previous study, Woodruff et al. (2025), blood samples were collected immediately after treatment (“Immediate”). **B) Biomarkers of carryover effects of heat.** Boxplots show z-score normalized gene expression for heat-treated (orange) and control (blue) nestlings sampled 24 hours after the heat challenge ended (“Carryover”). Symbols denote sex (circles = females, triangles = males).

## Methods

### Study subjects and location

Our experiment targeted nestling Tree Swallows (*Tachycineta bicolor)* breeding at an array of nestboxes in Knoxville, TN, USA (35°54’09.8"N 83°57’22.9"W) in May-Jun 2023. We checked nests every few days to determine lay dates and then checked daily on the projected hatch date. Tree Swallows hatch asynchronously, and we assigned hatch date (‘D1’) as the day the majority of the nestlings hatched. On D8 or D9, we visited the nest again to habituate birds to the experimental setup, placing dummy heat packs along two nestbox walls and one iButton temperature logger. Each iButton was programmed to read every 10 minutes, and it was fastened onto the nest cup surface, adjacent to where chicks reside and facing down to measure the heat coming up from the heat packs that would be placed beneath the nest on the trial day.

We timed our experimental trials for D12 post-hatch, allowing ± one day (D11-D13) to ensure treatment counterbalance by date and therefore yielding an average timing of D12.2 ± 0.1. By this age, nestlings are fully endothermic (Marsh, 1980), and they have reached their adult-like asymptotic mass (McCarty, 2001). We focused on nestlings because young birds are easy to manipulate, they tend to have a narrower window of thermal tolerance than adults (McKechnie, 2021), and insights from this sort of vulnerable yet critical period of development may have especially strong predictive value for understanding heat impacts on wild populations (Andreasson et al., 2020; Pottier et al., 2022).

### Experimental heat challenge and sampling protocol

Our experimental design used air-activated heat packs (Uniheat 72hr, hereafter “packs”), which are a tractable method for field-based thermal manipulations of the nest (Albert et al., 2023; Woodruff et al., 2023). These packs consist of sawdust, iron, and vermiculite, and they emit heat once exposed to air. Using these packs, we generated sublethal heat for four hours, i.e. sufficient time for widespread transcriptional responses to occur. Based on previous implementations (Derryberry et al., in press; Woodruff et al., 2025a), we knew that three heat packs would achieve temperatures in the nest cup that were likely to exceed the upper end of the thermoneutral zone (∼37.5°C avg for small songbirds, extracted from Appendix S1 in Wolf et al., 2017), but unlikely to reach lethal limits that are thought to occur with prolonged exposure ≥ 45°C (Pollock et al., 2021). Therefore, we placed one heat pack under the nest, one pack along the back wall of the nestbox, and one pack along the wall opposite the side door. Controls received spent packs, placed in these same positions. To ensure warming at experimental nests, packs were opened ∼1 hour before their placement in the nest. Treatments were counterbalanced by time of day, with an average start time of 11:14 local time (range: 10:01 to 12:33). Treatments were also counterbalanced by brood size (4.7 ± 0.3 nestlings, range: 2 to 6 per nest). Each trial lasted four hours (4h01 ± 0h01).

At the end of the heat challenge, we removed all experimental materials from the nest. We also weighed chicks to the nearest 0.1g with a digital scale, and we banded each chick with a USGS aluminum band for individual identification. We also confirmed sublethality at this time; all nestlings survived the challenge, as anticipated.

Twenty-four hours later (23.9 h ± 3 min), we returned to sample blood from two nestlings per nest. We targeted two nestlings to ensure sufficient yield for downstream analyses, aiming for individuals of median size to avoid potential mass-related confounds of heat tolerance or sensitivity. Each blood sample was ∼50-100 uL, which is well below volumetric limits of 1% body mass (∼200 uL for average nestling mass of ∼20g). We collected blood as soon as we approached the nest, with samples snap frozen on dry ice immediately thereafter and transferred to -80 storage later that day. Before returning nestlings to their nest, we weighed chicks and again confirmed welfare; all nestlings remained alive. We later found that all control and heat-treated nests had chicks fledge, again confirming the sublethality of this experiment.

### Molecular methods and informatics

We extracted total RNA from whole blood using Trizol, following the manufacturer’s protocol. High quality RNA was shared with our genomics core facility (n=9 controls, n=10 heat-treated individuals, each from a separate nest). Library prep and sequencing followed protocols we have used in the past for whole blood transcriptomes in this species (Bentz et al., 2019; Woodruff et al., 2025a). Briefly, we used a TruSeq Stranded messenger RNA kit to generate mRNA libraries, which were sequenced via NextSeq 500 in 75-cycle paired-end reads. To get the best mapping, we enhanced annotation of the *Tachycineta bicolor* genome [GCA_025960845.1] (Tseng et al., 2023) with the *Hirundo rustica* genome [GCF_015227805.2] (Secomandi et al., 2023) by assigning scaffolds to *H. rustica* chromosomes with RagTag (Alonge et al., 2022) and then using Liftoff (Shumate and Salzberg, 2021) to annotate the Tree Swallow genome with 17,707 genes. Next, we used STAR (Dobin and Gingeras, 2015) to map our samples fastq files to this enhanced genome. Samples had between 6.28 and 33.53 million reads (12.91 ± 1.58) with between 73.63% and 83.10% unique mapped (79.23% ± 0.51).

To assign sex to each individual, we leveraged the fact that birds have little to no dosage compensation (Itoh et al., 2007; Julien et al., 2012), and so, for Z-linked genes, ZW individuals (females) should have approximately half of the expression compared to ZZ individuals (males). Therefore, following the methods of (Hansen et al., 2022), we used a high-quality chromosome-level genome from *H. rustica*, focusing on the sex chromosomes W and Z. We also analyzed chromosome 5 as a representative autosome because it has a similar number of total genes compared to the Z and therefore provides a useful normalization against which we compared number of reads from each sex chromosome. This creates a Z:autosome ratio and W:autosome ratio, which are commonly used for sex identification (Disteche, 2016). Among our samples, we observed a clear bimodality for both these ratios: for the Z:A ratio, one group clustered around a ratio of 0.61 ± 0.01 and another around 1.01 ± 0.01 (Figure S1), consistent with the literature (Uebbing et al., 2013). We validated that this clustering represented chromosomal sex with data from six individuals of known sex who had been euthanized in earlier work (Woodruff et al., 2025a), with 100% accuracy. Though we did not know the sex of sampled individuals in the field, this resulted in four relatively balanced groups: control female (n=3), control male (n=6), heat-treated female (n=5), heat-treated male (n=5).

### Carry-over effects on blood gene expression

To identify genes with differential expression (DEGs) on the day after the experimental heat challenge, we used DESeq2 (version 1.42.1) in R/Bioconductor (R version 4.3.1) (Love et al., 2014). First, we filtered genes to focus only on those with at least 5 reads in at least half of the samples. Additionally, we filtered out 14 globin genes *(DGB, CYGB, IGSF21, JUP, LOC120748414, LOC120748415, LOC120757773, LOC120759534, LOC120759626, LOC120759893, LOC120764989, MB, NGB, RHEX)* as these genes are exceptionally highly expressed in bird blood, accounting for 36-54% of total reads per sample. These genes therefore have a disproportionate impact on dispersion estimates, which can mask true biological variation. At the end of these filtering steps, we retained 9,304 genes for DESeq2.

Next, we used a series of models to quantify the effects of heat treatment and sex. For our primary approach to modeling carryover effects of heat, we used an *additive model* that controlled for sex (∼ Treatment + Sex). To test the effects of continuous variation in temperature on gene expression, we used a similar additive model (∼Nest temperature + Sex) with normalized iButton nest temperatures on the trial day. To identify genes with sex-dependent carry-over effects of heat, we also evaluated an *interaction* model (∼Treatment*Sex), using pairwise Wald tests to identify any sex-specific carry-over effects of heat. To directly compare the magnitude and direction of any sex-specific responses to heat, we used a rank-rank hypergeometric overlap analysis (RRHO2) (Cahill et al., 2018; Plaisier et al., 2010); this approach compares two rank-ordered lists of DEGs, in this case the female- and male-specific heat DEG from the interaction model.

To maximize biological inference from these models, we adjusted the FDR threshold for DEG calling to match the distribution of p-value data from each model. For each model, we calculated the percent variance explained for a given gene by each of the terms in the DEG model (e.g. Sex or Treatment), using the R package variancePartition (Hoffman and Schadt, 2016). In this manner, we could interpret the average impacts of sex or treatment across all genes. Additionally, to determine the impacts of sex or treatment on just autosomal genes, we created new DEG models without the sex chromosome genes and again calculated the percent variance per gene.

We used two approaches to infer function within key gene sets. First, for the hundreds of genes that were differentially expressed between the sexes, we evaluated overrepresentation of gene ontology (GO) terms using ClusterProfiler (Wu et al., 2021) and enrichplot (G, 2025). We focused on sex-biased DEGs that were not on a sex chromosome, analyzing male-biased and female-biased autosomal DEGs separately. Second, for the more moderate differential expression related to carryover effects of heat, we used Gene Set Enrichment Analysis (GSEA) because this approach identifies coordinated changes across many genes with shared functions without a pre-set p-value cutoff. Our GSEA ranked all genes based on their signed -log10 p-value associated with treatment in the additive DEG model. For both approaches, we created a functional GO term reference using AnnotationForge (Carlson and Pagès, 2026) with the *H. rustica* genome (NCBI Tax ID: 43150), and we used genecards to aid our interpretation of specific genes (Stelzer et al., 2016).

### Comparing carry-over effects to immediate effects of heat

To contextualize these carryover effects of acute heat, we compared results from this main study to that of a previous study in which transcriptomic data were collected from blood immediately after four hours of heat (Woodruff et al., 2025a); see ***Figure 1A***. Briefly, that study used the same heat-pack methods, elevating nest temperatures by 4.5°C for 12-day-old Tree Swallow nestlings; RNA-seq showed differential expression for genes related to antioxidant defenses, inflammation, metabolism, ubiquitination, and especially heat shock proteins.

To generate a robust comparison between this previous heat challenge and the current experiment, we needed identically derived model outputs from the same bioinformatic pipeline. Therefore, we returned to the raw sequence files from this published project and performed the same mapping, quantification, filtering, and sexing as above (Figure S2). Samples had between 18.14 and 30.67 million reads (25.19 ± 2.10) with between 85.86% and 88.65% uniquely mapped reads (87.22% ± 0.43). This yielded 10,240 genes for DEG analysis, which we ran through the same additive DEG model (∼Treatment + Sex). Critically, we again used RRHO2 to contrast these immediately heat-reactive genes to the heat-reactive genes identified in our carryover effects experiment. In this case, RRHO2 compares lists of genes based on the magnitude and direction of their heat-associated log2-foldchange, identifying genes that are enriched for especially concordant (similar) or discordant (different) responses to heat at the two timepoints.

## Results

### Treatment effects on nests and nestlings

Nest cup temperatures for heat-treated nests were significantly hotter than those of controls during the four-hour trial (*t*(16.95) = -6.67, *p* < 0.001). Control nests were 35.1°C ± 0.6°C, whereas hot nests were 40.3 °C ± 0.6°C, for an average treatment-induced elevation of 5.2°C. On the subsequent day, when no thermal manipulation occurred, treatments had similar temperatures in the nest cup (*t*(16.57) = -0.52, *p =* 0.61): 33.7 ± 1.5°C in controls and 34.7 ± 1.3°C in experimental nests. Treatments did not significantly differ in body mass (*t*(11.81) = 0.88, *p =* 0.39), nor did they differ in change in body mass from the first to second day of sampling (*t*(11.06) = 0.74, *p =* 0.48). On both days, nestlings averaged 20.3g ± 0.5, as is typical at this age when nestlings invest most growth into feather production instead of mass gain.

### Sex differences in gene expression

Sex explained marked variation in blood gene expression (Figure S3). Specifically, our additive DESeq model showed that, on average, sex explained 11.7% and treatment explained 4.7% of the variation in gene expression (Figure S4a). Globally in nestling blood, we identified 138 female-biased DEGs (33 on the W, 4 on the Z) and 306 male-biased DEGs (2 on the W, 245 on the Z; FDR < 0.01; Figure S5). Excluding genes on the sex chromosome, we still found that the average percentage of variation due to sex was 9.6% and treatment was 4.8% (Figure S4b), suggesting that sex differences are not simply an artifact of chromosome copy number variation. Among these autosomal genes with sexually dimorphic expression, the male-biased genes were not enriched for any particular function in GO overrepresentation analysis. However, female-biased autosomal genes were enriched for diverse biological processes and molecular functions, including mRNA metabolism, ubiquitin, and many GO terms related to protein refolding, including heat shock protein binding (Figure S6). Four of these female-biased DEGs are HSPs: *HSPD1, HSPA2, HSPD1, DNAJB4*.

### Carry-over heat effects on gene expression

Focused on the treatment effect, our additive DEG model identified 4 genes with significant upregulation 24-hours following a heat challenge (*LAMA3, TMEM181, ATP1B1, RASGEF1A*) and 2 genes with significant downregulation, relative to controls (*LOC120758741, LOC120752683*) (FDR < 0.16); ***Figure 1B***. Of these six heat-responsive DEGs, two were also correlated with subtle among-nest differences in temperature on the preceding day (*TMEM181, LOC12075683*; Figure S7). GSEA analysis of the overall transcriptomic carryover effects of heat showed enrichment related to *nuclear receptor activity* and *ligand-activated transcription factor activity* (Figure S8), meaning that genes with these functions show coordinated shifts on the day after a heat challenge. These genes include *RARA, THRA, RXRG, RARG, NR4A3,* and *RXRA*.

Using the output of our interaction model on sex-specific heat effects, we identified 91 heat-linked DEG in males and 58 DEG in females; 51 of these were shared between the sexes (FDR < 0.05; Figure S9). Consistent with this view, RRHO2 showed high concordance between males and females in how each gene responded to a heat challenge the day before (adjusted p-value < 0.05): 4,264 and 4,001 genes were significantly enriched for overlapping decrease and increase across sexes, respectively, and no genes were enriched for discordance (Figure S10).

### Comparing carryover effects to immediate effects of heat

To understand the temporal dynamics of transcriptomic response to heat challenge, we compared results from this main 24-hour study to a previous study in which blood data were collected at the end of a four-hour heat challenge (Woodruff, Tsueda, et al., 2025). We reconstructed those results, now accounting for the previously unknown effect of sex, with 67 upregulated DEGs in the immediate aftermath of heat and 18 downregulated DEGs (FDR < 0.001; (Figure S2, S11, S12, S13). Our direct comparison of these immediate vs. carryover effects of heat revealed marked temporal dynamism. Specifically, RRHO2 identified only 12 genes enriched for concordance (i.e. similar) patterns at the two time points (***Figure 2A***), half of which showed a heat-associated increase in both experiments (*ARHGEF38, BPGM, MARCHF3, CHST2, GNPDA2, ENPP4*) and half of which showed a heat-associated decrease in both experiments (*ST7, APRT, MRM3, PIK3CD, NR4A3, ACP6;* ***Figure 2B***). None of these concordant genes were DEGs in either within-experiment analysis. More importantly, an additional 4,668 genes were significantly discordant in their response to heat at the two timepoints (***Figure 2A***). These discordant genes included many of the DEGs from both studies, including 4 from our main carryover effects analysis and 61 from the previous immediate heat response analysis. For the 56 HSP genes expressed in blood in both projects, 23 are discordantly expressed at the two timepoints, including six of the eight HSP DEGs seen at the end of the four-hour heat challenge (Figure S13).

**Figure 2.**
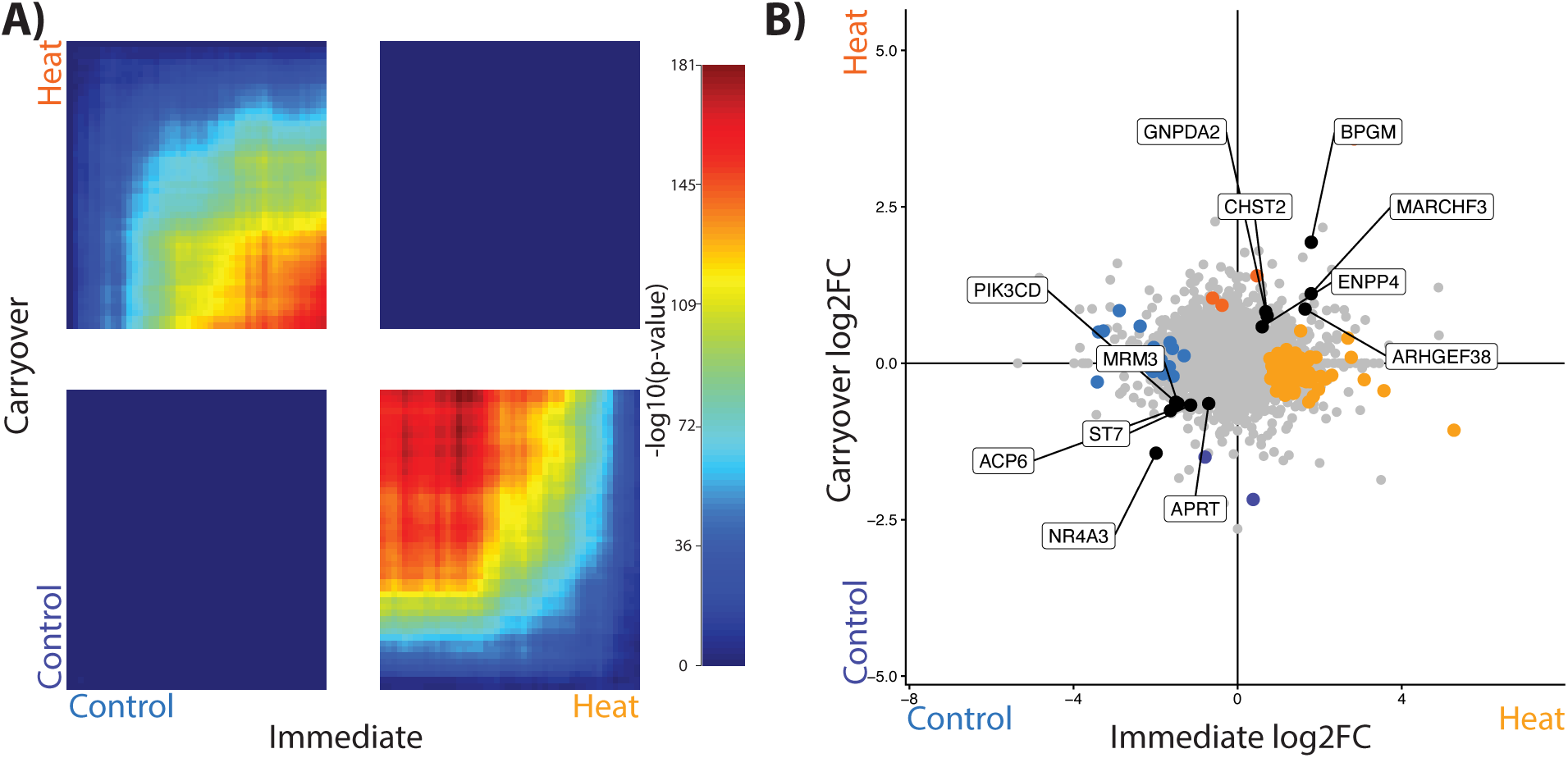
Comparing Immediate and Carryover Effects of Heat. **A)** Immediate and Carryover Effects show marked discordance, based on rank-rank hypergeometric overlap test. The y-axis is a ranked list of genes, based on the significance and direction of the carryover effect of heat on expression. The x-axis is a separately ranked list of genes, based on the significance and direction of the immediate effect of heat on expression. Each pixel represents the overlap between corresponding bins of genes from the two ranked lists, with color indicating the significance of overlap (adjusted hypergeometric –log(p-value)). Genes in the lower-right and upper-left quadrants show opposing response to heat at the two different sampling points, whereas genes in the upper-right and lower-left quadrants show similar responses at the two sampling points. **B)** Scatterplot of these same 9,184 genes, contrasting the heat-associated log2 foldchange (log2FC) at the immediate sampling point (x-axis) versus the carryover sampling point (y-axis). Visualized this way, we see the smaller subset of genes that are enriched for concordance between experiments (colored black and labeled). DEGs from each sampling point are also colored to indicate the direction of differential expression, with warmer colors representing higher expression in heat-treated nestlings and cooler colors representing higher expression in control nestlings. Lighter colors denote immediate effects DEGs (yellow, blue) and darker colors denote carryover effects DEGs (orange, indigo).

## Discussion

As temperatures rise and many bird populations decline, ornithology has both a basic and applied need for biomarkers of heat that can be deployed in the field. Via our analyses of global gene expression in a drop of bird blood, we identify several genes whose transcript abundance tracks prior sublethal heat, with differential gene expression at least 24 hours after the stressor subsided. Notably, these carryover effects on gene expression were distinct from genes that responded to heat in its immediate aftermath, highlighting the temporal complexity of biomarker validation. We also show that – despite pronounced sexual dimorphism in baseline gene expression in nestling Tree Swallows – the transcriptomic response to heat is strikingly similar in males and females, supporting the utility of these biomarkers across individuals, even in wild outbred populations with naturally diverse environmental and genetic backgrounds. Through these findings, we provide a minimally invasive and temporally calibrated diagnostic for studying climate change in free-living birds, alongside a roadmap developing similar tools in other study systems.

### Biomarker validation

For biomarkers of heat to be informative, it is essential to define whether they reflect recent heat exposure or the physiological responses that unfold over time after that heat exposure. Our analysis of carryover effects of heat on the blood transcriptome reveals very little overlap with the initial transcriptomic signatures of heat. For instance, heat shock protein mRNA or protein abundance have been used across diverse organisms to detect exposure to heat (Etches et al., 2008; Kumar et al., 2003; Sejian et al., 2018). Because HSPs are molecular chaperones, their rapid upregulation induced by heat or other stressors is thought to help protect or repair proteins and cells from heat-associated damage (Feder and Hofmann, 1999). However, our results show that the initial HSP response has statistically returned to control-like levels within 24 hours after a heat event (Figure S13), suggesting HSPs may have utility as biomarkers only within a relatively narrow window during or immediately after a heat event. Instead, our gene set enrichment analysis points to a coordinated carryover effect of heat on nuclear receptor signals, with key examples related to lower expression of thyroid hormone (*THRA*), the glucocorticoid-associated orphan receptor (*NR4A3*) (Srikanth et al., 2019), and several retinoid receptors (RXRG, RXRA, RARA, RARG). Retinoid receptors are known to be thermally unstable (Yang et al., 2021) and they require HSP90 for normal functioning (Holley and Yamamoto, 1995). During a heat event, these unstable nuclear receptors may be impaired while HSPs are tied up with other repair duties, and therefore the carryover effect may explain why these functional pathways have still not fully recovered a day later. More broadly, the temporally distinct waves of transcriptional activity that we see immediately and 24 hours after heat are consistent with similar lability in the brain after a social challenge, when different genes of distinct molecular functions respond earlier versus later in a post-stimulus cascade (Bukhari et al., 2017; Saul et al., 2019; Shpigler et al., 2017).

Amid this temporal dynamism, a subset of genes exhibits consistent responses to heat across timepoints, making promising candidates for biomarker development. Such stability is particularly valuable for researchers who are not present to observe heat exposure directly but instead seek to infer thermal history from point sampling. Among the 12 genes we identified with consistent upregulation or consistent downregulation at our two sampling points, a few bear mention due to their effect size (log2fold-change ≥ 1 in both contrasts) and likely function. For instance, the consistently upregulated gene *BPGM* encodes an enzyme that decreases oxygen affinity in hemoglobin, and *MARCHF3* encodes a ubiquitin ligase that marks damaged proteins for disposal. *NR4A3*, on the other hand, was consistently downregulated, and it encodes a transcription factor with connections to both glucocorticoid signaling and heat (Srikanth et al., 2019). Notably, though, none of these candidates were DEGs at either experimental timepoint, and we urge further inquiry before they are considered viable, lasting markers of heat.

We see the greatest promise for six specific genes as biomarkers of the carryover effects of heat. *LAMA3, TMEM181, ATP1B1*, and *RASGEF1A* are upregulated 24 hours after a heat stress event, whereas *LOC120758741* and *LOC120752683* are downregulated (**Figure 1B**). These downregulated genes are not well annotated, but they are related to nucleoside transport and cytokine signaling, respectively. For the four genes with upregulated expression after heat, most have been used as biomarkers in clinical settings (Islam et al., 2023; Wang et al., 2024), and all have some connection to stress: *LAMA3* encodes a component of the extracellular matrix that affects red blood cell adhesion under oxidative stress (Maria Alejandra et al., 2021). This gene also exhibits changes in its methylation after exertional heat stroke in mice (Tijjani et al., 2019) and its sequence exhibits signatures of selection in heat-adapted West African cattle (Cao et al., 2021; Huang et al., 2025). *ATP1B1* encodes part of an ion pump that is known to interact with HSPs (Zheng et al., 2024); it has also been linked to thermal adaptation in poultry breeds (An et al., 2025; Lassiter et al., 2025; Nawaz et al., 2026) and heat tolerance in experimentally heat stressed broilers (Celesnik et al., 2024; Emilsson et al., 2018; He et al., 2023; Katoh and Katoh, 2006; Xu et al., 2023). *RASGEF1A* and *TMEM181*, which are both involved in cellular signaling cascades, have published connections with resilience to radiation stress in pigs (Chopra et al., 2020) and toxin exposure in humans (Correia et al., 2022), respectively. Critically, *TMEM181* and *LOC120752683* also track the exact temperature in the nest cup the previous day (Figure S7), meaning they may provide information on nestling microclimates even when temperatures themselves cannot be measured.

### Sex-specific considerations

One particularly striking element of our results is the massive sexual dimorphism in the blood transcriptome, despite sampling altricial nestlings with no discernable external sexual traits, still 10 months before their gonads fully develop. Since birds do not have global dosage compensation, some degree of dimorphic gene expression is expected, particularly for genes on the Z or W sex chromosomes (Itoh et al., 2007; Julien et al., 2012), a result we recapitulated here (Figure S1; S5). This basic biology allows for sex identification via gene expression of W-linked genes, like *SMAD4, RBMX, NKRF*, or *UBE2A* in our study. Though DNA-based sexing via PCR has negated the need for laparotomies, researchers sometimes must choose between DNA or RNA applications, given limited blood volumes and the cost of extraction. Our findings introduce new candidates for RNA-based biomarkers of sex, which work with as little as 25-100uL of blood. Future testing could convert these candidates into semi-quantitative gel-based PCR, instead of more costly or time-consuming RNA-seq or quantitative PCR. Validation across species is also key, considering that the sex ratio of gene expression can evolve rapidly among species (Miller-Crews et al., 2025), though the high degree of chromosomal synteny across bird taxa suggests high potential for broad application (Ellegren, 2010).

Importantly, 36% of sex-biased genes in our dataset are in fact autosomal (Figure S5). This result is strikingly similar to the degree of sex bias seen across the transcriptome of a sexually dimorphic and socially relevant brain region in adult Tree Swallows (Miller-Crews et al., 2025), potentially implying an element of functional sex differences in the blood too. Though this interpretation requires further validation, it aligns with similar findings in adult human blood (Jansen et al., 2014) as well as ornithology’s recent interest in the functional activity of blood cells as modulators of other elements of the phenotype (Loveland et al., 2025). In our analysis, female-biased autosomal genes are enriched for GO categories related to response to unfolded proteins and *heat shock protein binding*, the main function of HSP genes. In fact, HSPs were enriched among the DEGs for sex with 4 expressed more highly in female blood than male blood. Taken together with similar sex differences in HSPs in humans and in poultry (Balakrishnan et al., 2023; Voss et al., 2003), this suggests that the sexes may have differential readiness to respond to heat when it arises.

Layered atop these main effects of sex, we saw overwhelmingly similar heat responses in females and in males. 53% of the heat-associated DEGs within each sex overlapped with that of the other sex (Figure S9, S10), and even among those that were not DEG, the sexes still showed high concordance in the direction and magnitude of heat effects across the transcriptome (Figure S10). More narrowly, 154 genes showed evidence of sex-modulated response to heat, with one sex responding more strongly than the other. This pattern is consistent with sex differences in sensitivity to or tolerance of other anthropogenic stressors (Dantzer et al., 2014). More broadly, we do not yet know whether the early postnatal differences we see here in the blood persist into adulthood, and our sample size for detecting sex-specific effects remains modest. Nonetheless, incorporating sex into gene expression analyses can uncover potentially relevant variation that may otherwise be masked.

### Conclusions and future application

Our collective results uncover promising biomarkers for understanding heat impacts on birds, including genes whose carryover effects of heat transcend inherent sex differences in expression. That said, one key assumption implicit to biomarker application is the notion that variation in the marker reliably maps to some element of prior stress, be it exposure, response, or recovery. Though we connected several genes to temperature variation, biomarker mapping is rarely straightforward for several reasons (Dantzer et al., 2014; Johnstone et al., 2017). Of relevance here, for example, altricial nestmates move through different microclimates within the nestbox (Derryberry et al., in press), and siblings exhibit correlated but non-identical gene expression responses to heat (Woodruff et al., 2025a). Individuals also differ in panting (Dawson, 1982), and these thermoregulatory behaviors may drive differences in the downstream physiological effects of heat (Lipshutz et al., 2022; Rosvall and Derryberry, in press). Early life heat exposure can reshape how birds respond to later heat stress (Hoffman et al., 2024; Hoffman et al., 2026), and coupled with the potential for local adaptation, populations may differ constitutively in gene expression even before a heat stress event (Woodruff et al., 2025b; Woodruff et al., 2022). Via repeated measures within individuals and comparisons among populations with distinct thermal regimes, the putative biomarkers we identified can be further calibrated against both short-term exposure and longer-term thermal history.

In conclusion, our research on nestling Tree Swallows uncovers both immediate and carryover effects of sublethal heat, which we hope to see explored in other systems. As the field continues to connect these and others putative biomarkers with performance (Woodruff et al., 2025a) (Svensson et al., 2024), we will gain insight on the valence of climate change’s lingering effects on birds, providing critical data needed to improve predictions for the future (Briscoe et al., 2023; Comte et al., 2024).

## Acknowledgments

For support in the field, we thank L Para, R Simberloff, M Lefeuvre, and the University of Tennessee (UT) ETREC Plant Science Unit. For support in the lab, we thank T Empson, D Rusch, A Buechlein, and Indiana University’s Center for Genomics and Bioinformatics. We also thank MJ Woodruff for sparking our interest in blood-based transcriptomic biomarkers of heat.

## Ethics statement

All research adhered to local, federal, and university regulations, including Tennessee IACUC #2709-0126, Federal banding permit 23900, and Tennessee Scientific Collection Permit 2859.

## Author contributions

EPD and KAR conceived the experiment and secured funding. EPD performed the main experiment and collected field data. IMC performed all bioinformatics and statistical analyses. All authors interpreted results. All authors contributed to the initial manuscript draft and ensuing revisions.

## Data Depository

Sequence data for both this experiment and previous experiment are on NCBI SRA under BioProject PRJNA1473279. All code for analysis and figures is available on github (https://github.com/imillercrews/TRES_heat_biomarkers_bl).

