## Supplemental for "Toward temporally calibrated biomarkers of heat stress in free-living songbirds"

A)

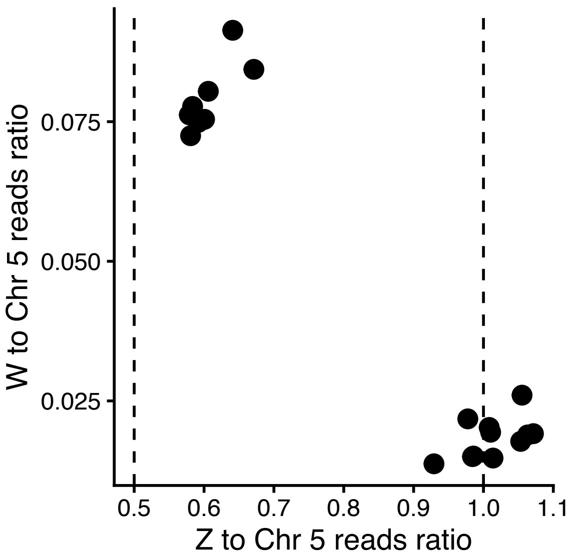


### Figure S1.

**A)** We determine sex for each of the samples in our main carryover effects study by comparing the number of reads for genes on the Z chromosome to those on similarly sized chromosome 5 (x-axis) and by comparison the number of reads for genes on the W chromosome and chromosome 5 (y-axis). Dashed lines indicate predicted theoretical Z to chromosome 5 ratio for ZZ males (1) and ZW females (0.5).

**A)**

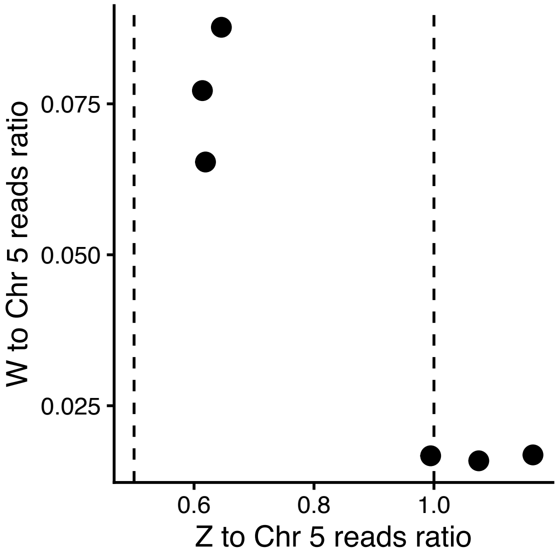


### Figure S2.

**A)** We replicated the same mapping, quantification, filtering and sexing protocols for the previously published immediate transcriptomic effects of heat, described in Woodruff et al. (2025). This included the removal of the same globin genes, which accounted for 16-28% of total reads. We again determined sex by comparing the number of reads for genes on the Z chromosome to those on similarly sized chromosome 5 (x-axis) and by comparison the number of reads for genes on the W chromosome and chromosome 5 (y-axis). Dashed lines indicate predicted theoretical Z to chromosome 5 ratio for ZZ males (1) and ZW females (0.5).

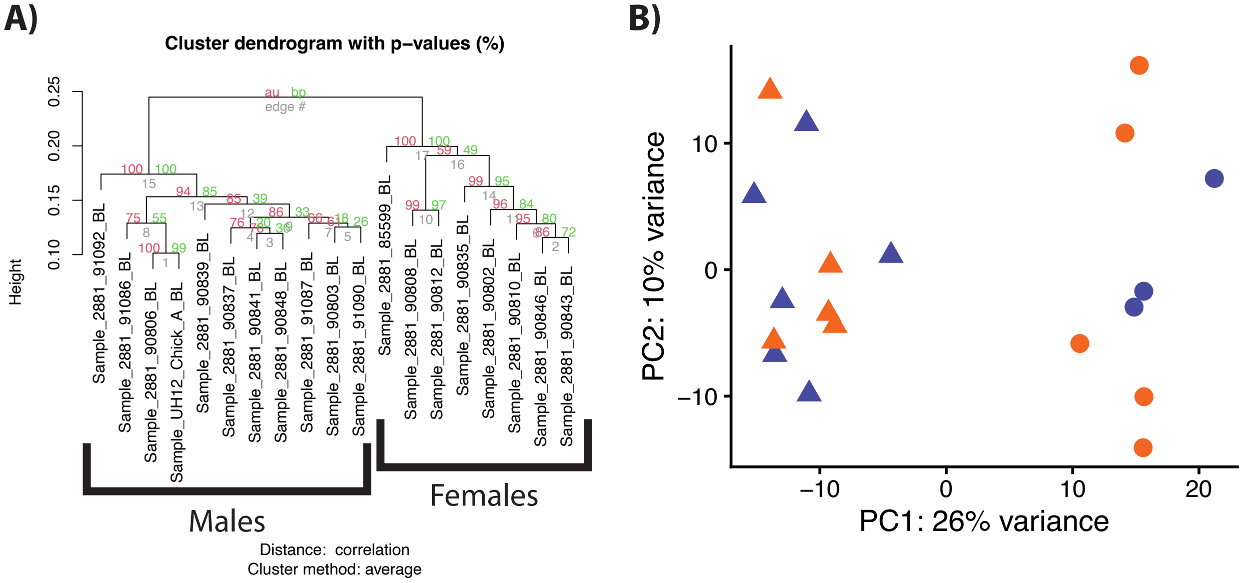


### Figure S3.

**A)** Hierarchical clustering shows clear separation between males and females, using normalized read counts from the 1000 most variable genes.

**B)** Principal component analysis of these data shows clear separation between males and females. The top ten loadings for PC1 including 7 W genes, 2 Z genes, and 1 autosomal gene, which all load positively towards females. Among these, the top-loading gene is the W gene *SMAD4*. Shape indicates sex (males = triangle, females = circles) and color indicates treatment (heat = orange, control = indigo).

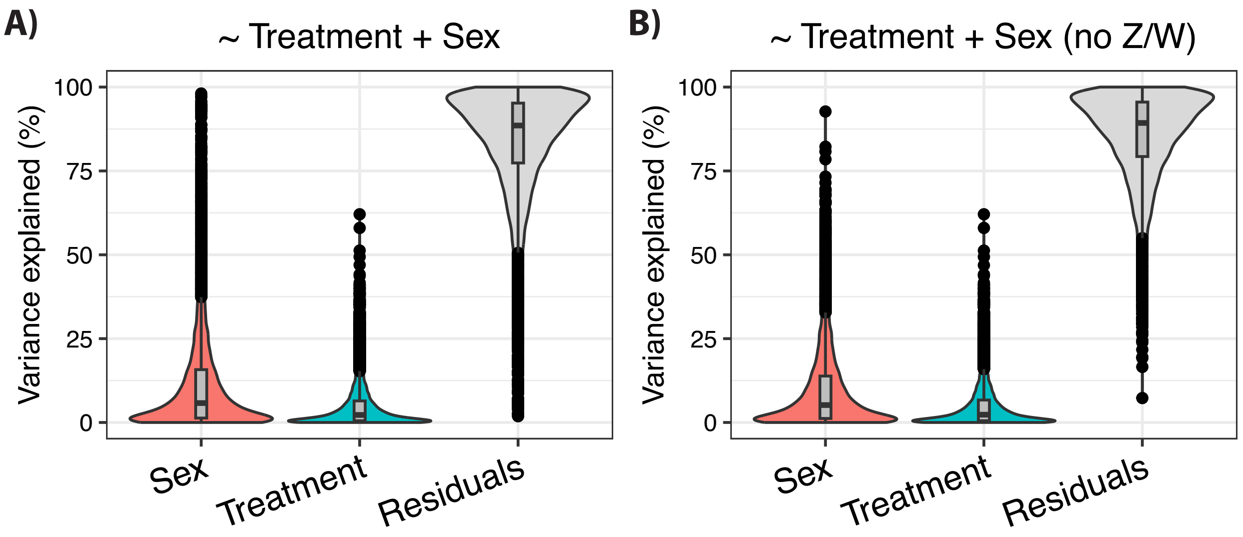


### Figure S4.

**A)** Across all genes, the percent of variance explained by each main effect in the additive DEG model, summarized with a boxplot. On average, sex explained 11.7% and treatment explained 4.7% of the variation in gene expression. Each dot represents one gene.

**B)** Across only autosomal genes, the percent of variance explained by each main effect in the additive DEG model, summarized with a boxplot. On average for autosomes, sex explained 9.6% and treatment explained 4.8% of the variation in gene expression. Each dot represents one gene.

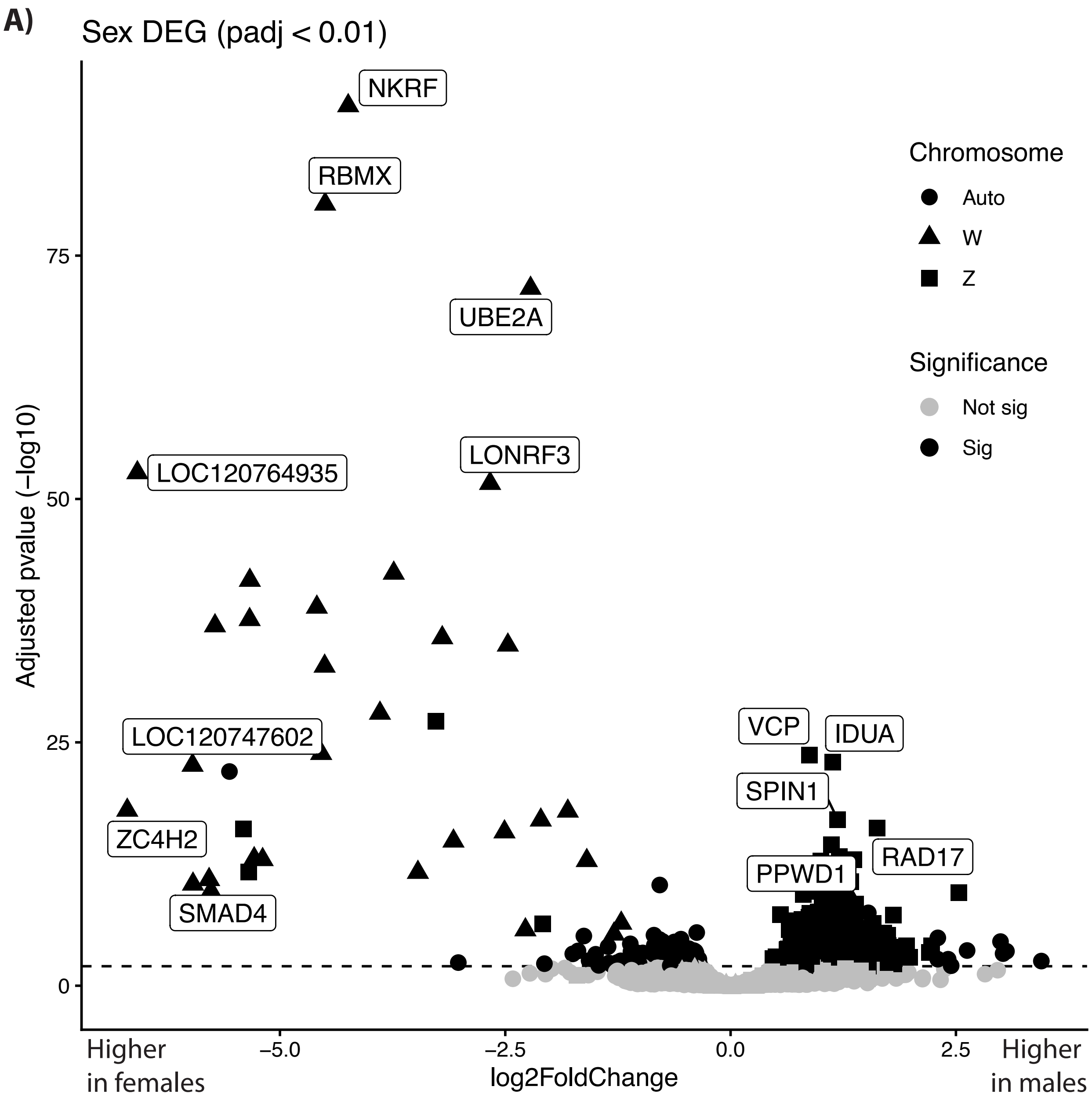


### Figure S5.

**A)** Volcano plot of significant sex-biased DEGs (FDR < 0.01; black points) across chromosomes (shape), seen in our main carryover effects experiment. Dashed horizontal lined denotes FDR significance threshold.

**A)**

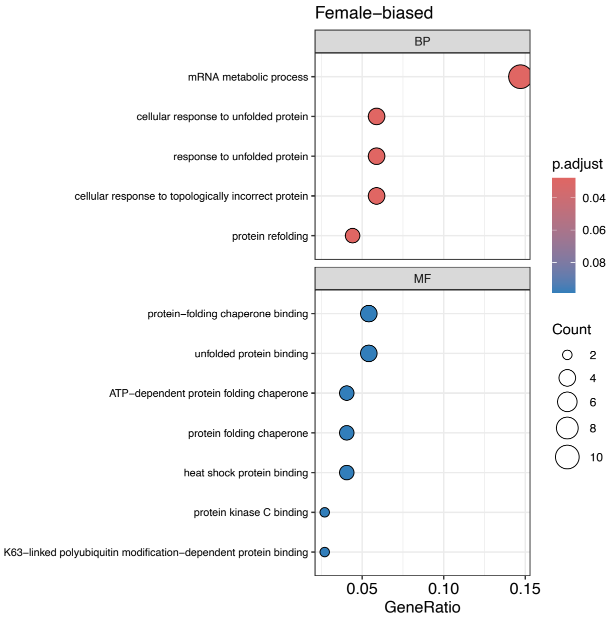


### Figure S6.

**A)** GO overrepresentation identified several enriched terms among female-biased autosomal DEGs, based on our additive model. Rows indicate significantly enriched GO terms among these leading-edge genes, with dots indicating gene ratio [proportion of GO term associated genes in list of significant genes] (x-axis), adjusted p-value (color), and number of genes enriched per category (dot size). Top label indicates GO category (BP = biological process; MF = Molecular function).

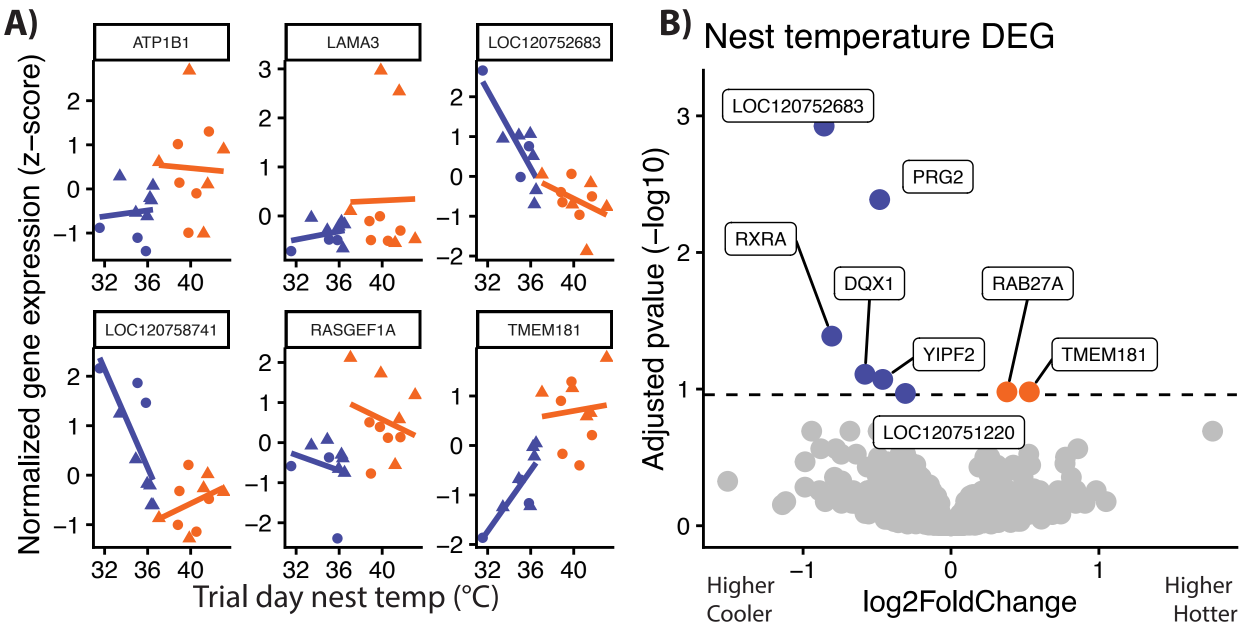


### Figure S7.

**A)** For the six main DEG linked to carryover effects of heat (shown in Figure 1b), we also plot their relationship with nest cup temperatures measured during the trial 24 hours before blood was sampled. Points indicate individuals with color denoting treatment (heat-treated = orange, control = indigo) and shape denoting sex (male = triangle, female = circle).

**B)** Volcano plot of significant DEGs (FDR < 0.11) when nest temperature is modelled as continuous variable, controlling for sex. Orange = genes whose expression is positively correlated with nest cup temperatures measured during the trial 24 hours earlier. Indigo = genes whose expression is negatively correlated with nest cup temperatures measured during the trial 24 hours earlier. Dashed horizontal lined denotes FDR significance threshold.

**A)**

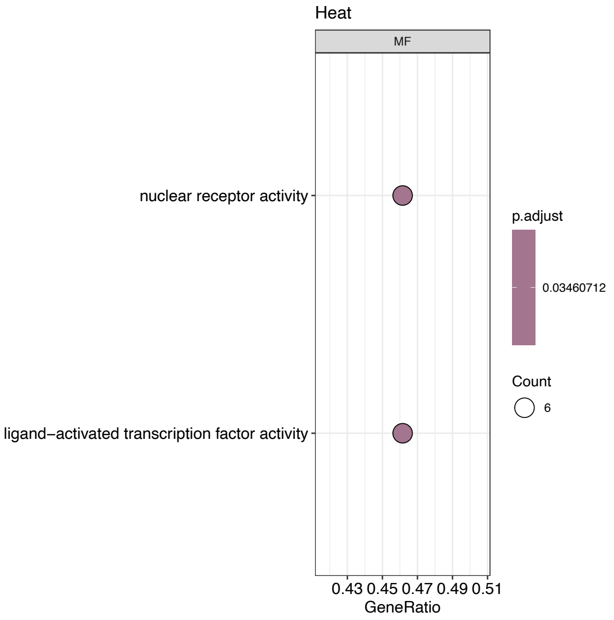


### Figure S8.

**A)** Gene Set Enrichment Analysis (GSEA) identified coordinated changes across genes with shared functions, based on our additive model of the carryover effects of heat. Rows indicate significantly enriched GO terms among these leading-edge genes, with dots indicating gene ratio [proportion of GO term associated genes enriched among all gene associated with that GO term ] (x-axis), adjusted p-value (color), and number of genes enriched per category (dot size). Top label indicates GO category (BP = biological process; MF = Molecular function).

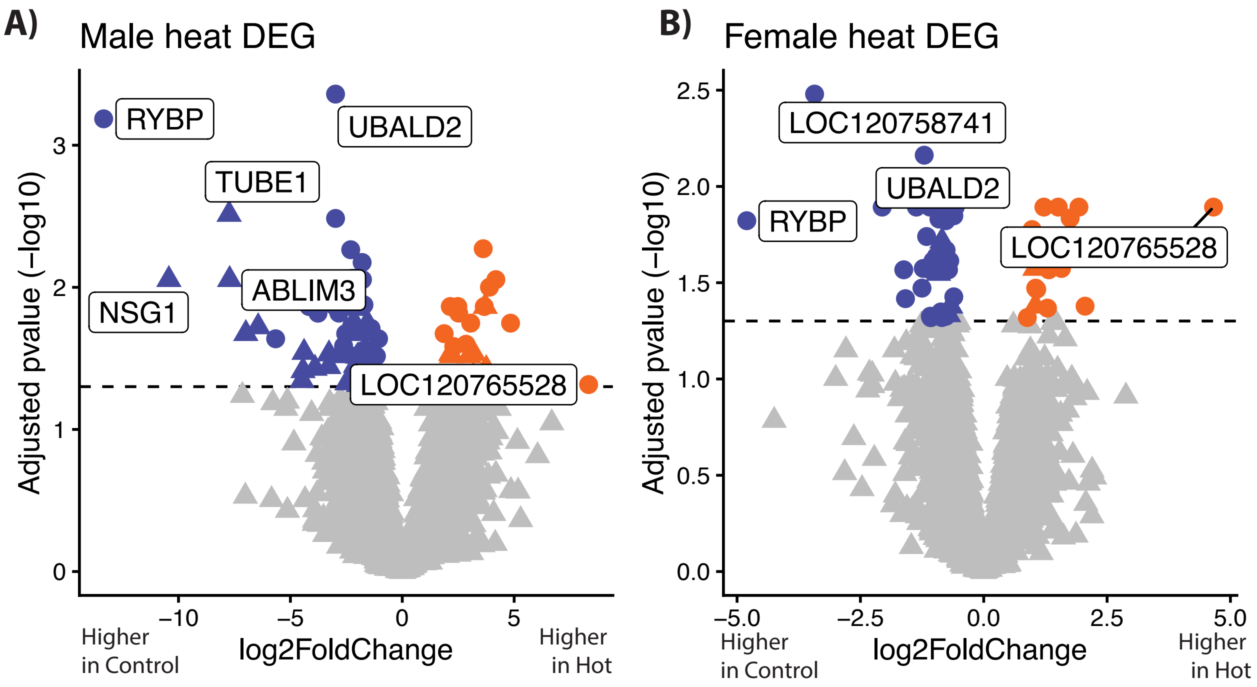


### Figure S9.

Volcano plot of significant DEGs by treatment in **A)** males and **B)** females, identified in our interaction model from the main carryover effects experiment. Post-hoc Wald tests showed that males have a significant increase in 22 genes and decrease in 69 genes, while females have a significant increase in 16 genes and decrease in 42 genes (FDR < 0.05). Color denotes for higher expression after heat-treatment (orange) or higher expression among controls (indigo). Dashed horizontal lined denotes FDR significance threshold.

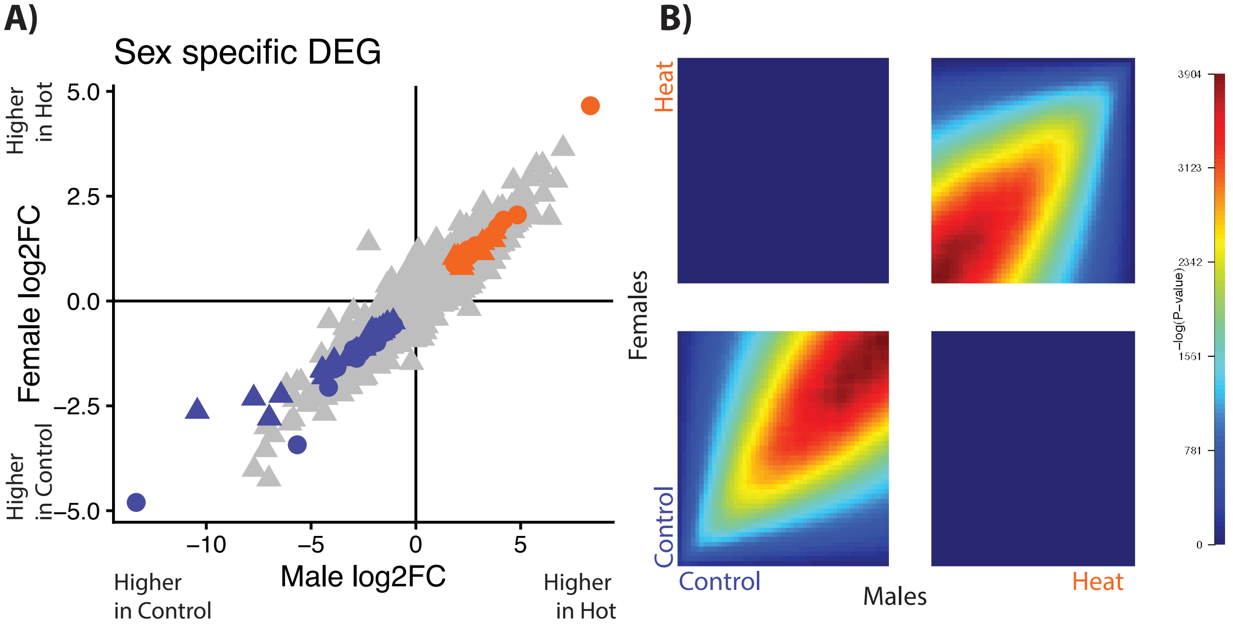


### Figure S10.

Females and males show overwhelmingly similar transcriptomic carryover effects of heat in the blood.

**A)** For each gene, we plot the heat-associated log2 foldchange (log2FC) in males (x-axis) to the log2FC in females (y-axis). Colored points indicating genes with significant carryover effects of heat in both the male and female analysis, with orange denoting higher expression after heat and indigo denoting higher expression in controls. Among these concordant heat-responsive genes, the strongest heat-associated upregulation was *LOC120765528* and strongest downregulation was *RYBP*.

**B)** Rank-rank hypergeometric overlap (RRHO) analysis statistically demonstrates high concordance between female carryover effects of heat and male carryover effects of heat. Control and Heat labels indicate if a gene is ranked higher in each treatment for each sex, respectively, based on its signed p-value. Each pixel in the heat map represents an overlapping bin of genes between the two rank ordered lists from the two sexes, with colors reflecting adjusted hypergeometric –log(p-value). Colored pixels in the lower-right and upper-left represent bins significantly enriched for concordance among the gene lists (e.g. increased expression in both males and females following heat challenge), while the remaining quadrants of lower-right and upper-left represent discordance.

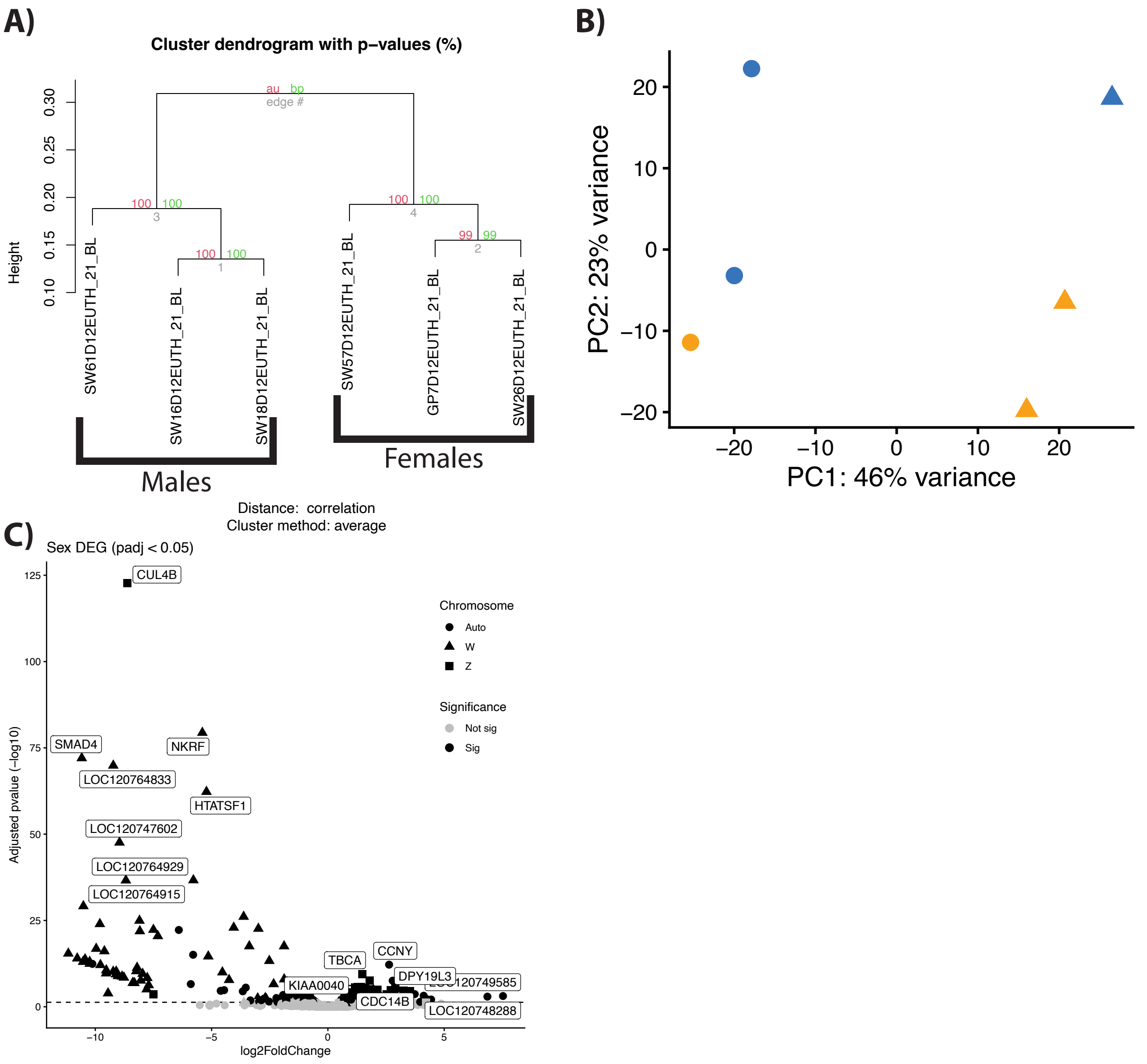


### Figure S11.

We replicated our same mapping, annotation, and QC to determine sex in samples from Woodruff et al. (2025), which measured gene expression at the end of a four-hour heat challenge.

**A)** Hierarchical clustering again shows clear separation between males and females, using normalized read counts from the 1000 most variable genes.

**B)** Principal component analysis of these data again shows clear separation between males and females. Shape indicates sex (males = triangle, females = circles) and color indicates treatment (heat = orange, control = indigo).

**C)** Volcano plot of significant sex-biased DEGs (FDR < 0.05; black points) across chromosomes (shape). We identified 324 female-biased DEGs, 57 of which are on the W and 3 on the Z, and we identified 326 male-biased DEGs, of which 2 are on the W and 182 on the Z (FDR < 0.05).

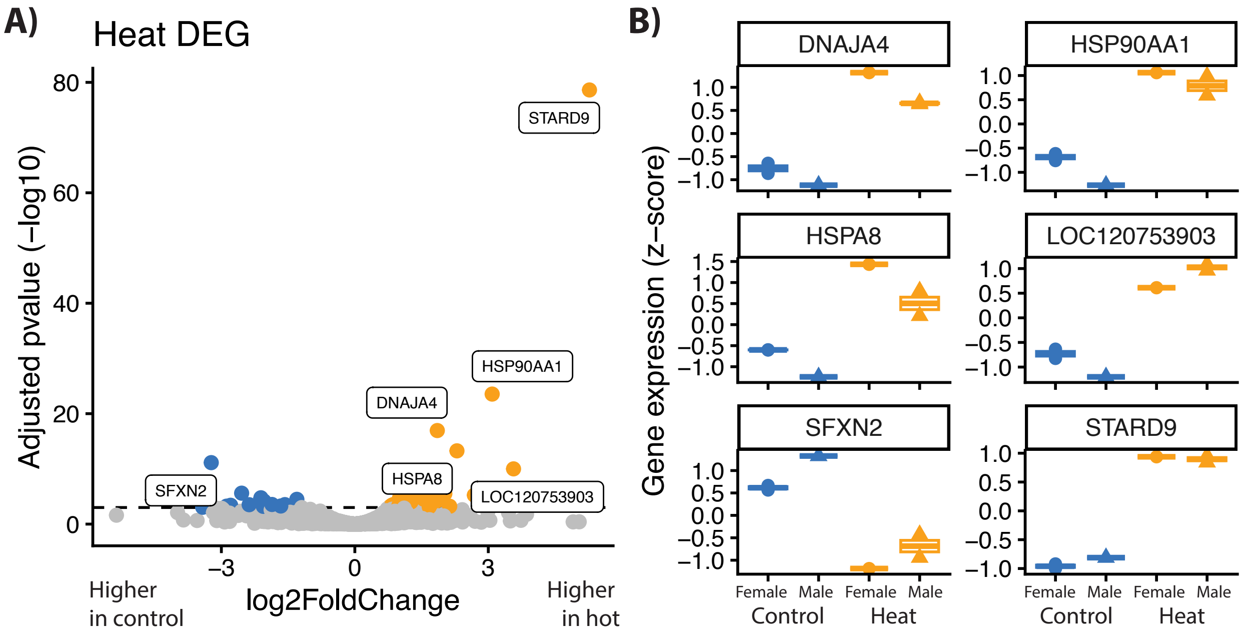


### Figure S12.

Replicating Woodruff et al (2025), but now accounting for the effect of sex, we again find the same DEGs in the immediate aftermath of a four-hour heat challenge.

**(A)** Volcano plot of significant (FDR < 0.001) DEGs by treatment. Yellow denotes genes with higher expression just after heat, whereas blue denotes higher expression in controls. The top upregulated genes are *STARD9, LOC120753903* and 8 HSP genes: *DNAJA1, DNAJA4, DNAJB4, DNAJC12, HSP90AA1, HSPA4L, HSPA8, HSPH1*. The top downregulated genes are *SFXN2, DPY19L3*. Dashed horizontal lined denotes FDR significance threshold.

**B)** A subset of the top heat-associated genes from Woodruff et al.'s data, plotting Z-score normalized gene expression based on treatment (heat-treated = yellow, control = blue) and sex (triangle = male, circle = female).

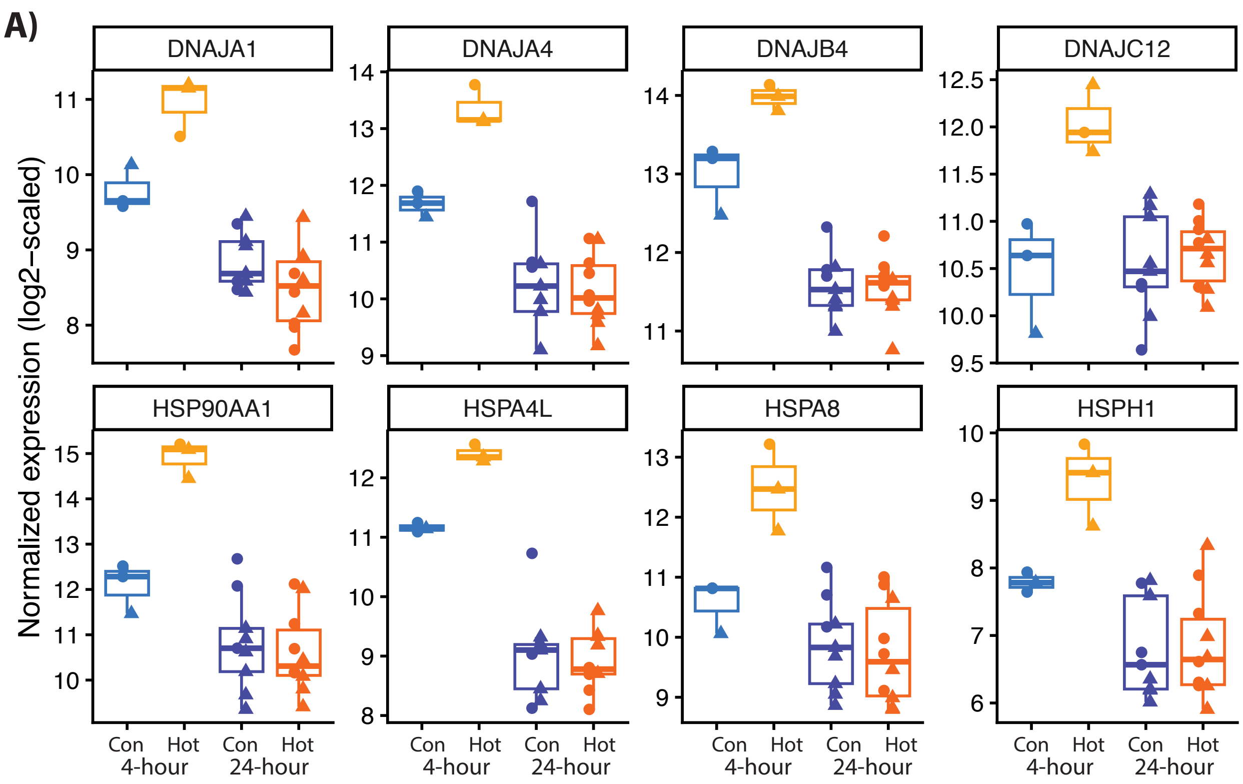


### Figure S13.

**A)** Among the 56 HSP genes expressed in blood in both projects, most show different heat associated responses in the immediate aftermath of heat (Woodruff et al. 2025) versus 24-hours later (our main study). We illustrate this point for the eight HSPs that were DEG at the end of the four-hours heat challenge, showing the Z-scaled normalized gene expression (y-axis) for each individual across treatment (immediate effects experiment: heat-treated = yellow, control = blue; carryover effects experiment: heat-treated = orange, control = indigo). Shapes denote sex (male = triangle, female = circle).
